# Immune Pressure is Key to Understanding Observed Patterns of Respiratory Virus Evolution in Prolonged Infections

**DOI:** 10.1101/2025.04.28.651059

**Authors:** Amber Coats, Yintong Rita Wang, Katia Koelle

## Abstract

For a number of viral pathogens, prolonged infections have been hypothesized to be the source of new variants that emerge and spread at the level of the host population. This potential role of prolonged infections has been highlighted most recently in the context of SARS-CoV-2, with variants of concern Alpha and Omicron likely to have evolved in immunocompromised individuals experiencing long-term viral infections. Analyses of sequenced viral samples from prolonged infections have indicated that there are several consistent patterns of evolution observed across these infections. These patterns include accelerated rates of nonsynonymous substitution, viral genetic diversification into distinct lineages, parallel substitutions across infected individuals, and heterogeneity in rates of antigenic evolution. Here, we use within-host model simulations to explore the drivers of these evolutionary patterns. Our simulations build on a tunably rugged fitness landscape model to first assess the role that mutations that impact only viral replicative fitness have in driving these patterns. They then further incorporate pleiotropic sites that jointly impact replicative fitness and antigenicity to assess the role that immune pressure has on these patterns. Through simulation, we find that the empirically observed patterns of viral evolution in prolonged infections cannot be robustly explained by viral populations evolving on replicative fitness landscapes alone. Instead, we find that immune pressure is needed to consistently reproduce the observed patterns. While our simulation models were designed to shed light on drivers of viral evolution in prolonged infections with respiratory viruses that generally cause acute infection, their structure can be used to better understand viral evolution in other acutely-infecting viruses such as noroviruses that can cause prolonged infection as well as viruses such as HIV that are known to chronically infect.

## Introduction

Many respiratory viruses, including influenza viruses and coronaviruses, typically cause acute infections that last less than two weeks (Einav *et al*., 2020; Bar-On et al., 2020). However, particularly in individuals who are immunocompromised or immunosuppressed, these viral infections can persist for much longer (Memoli *et al*., 2014; Baang et al., 2021; Dioverti *et al*., 2022; Machkovech et al., 2024). Sequencing of respiratory tract samples from individuals experiencing prolonged viral infections has revealed that a considerable amount of viral evolution can occur in these infections (Rogers *et al*., 2015; Xue et al., 2017; Kemp *et al*., 2021; Borges *et al*., 2021; Chen *et al*., 2021; Harari *et al*., 2022; Scherer *et al*., 2022; Khatamzas et al., 2022; Sonnleitner et al., 2022; Gonzalez-Reiche *et al*., 2023; Nussenblatt et al., 2022; Quaranta et al., 2022; Ko et al., 2022; Chaguza *et al*., 2023). Understanding these viral evolutionary dynamics is important for several reasons. First, these evolutionary dynamics can reveal whether the infecting virus has evolved escape from a patient’s immune response or treatment regimen (McMinn *et al*., 1999; Rogers et al., 2015; Jensen et al., 2021; Kemp et al., 2021; Khatamzas et al., 2022; Scherer *et al*., 2022; Khosravi *et al*., 2023), and as such could help inform treatment strategies for the focal patient and more generally for individuals experiencing prolonged viral infections. Second, new viral variants that emerge at the host population level may originate from viruses that evolved in individuals experiencing prolonged infections, as has been discussed for SARS-CoV-2 (Ghafari *et al*., 2022; Hill et al., 2022; Berkhout and Herrera-Carrillo, 2022). Characterizing patterns of viral evolution within individuals with prolonged infections could therefore help in surveillance efforts at the level of the host population and efforts to anticipate phenotypes of forthcoming variants.

Many studies have described patterns of respiratory virus evolution within individuals with prolonged infections (Rocha *et al*., 1991; Rogers et al., 2015; Xue et al., 2017; Kemp *et al*., 2021; Chen *et al*., 2021; Borges *et al*., 2021; Khatamzas *et al*., 2022; Scherer *et al*., 2022; Sonnleitner et al., 2022; Wilkinson et al., 2022; Gonzalez-Reiche et al., 2023; Nussenblatt *et al*., 2022; Ko et al., 2022; Harari et al., 2022; Riddell et al., 2022; Quaranta *et al*., 2022; Chaguza et al., 2023). In these studies and others, four evolutionary patterns are frequently observed: (1) Nonsynonymous substitution rates tend to be higher than synonymous substitution rates (Choi *et al*., 2020; Harari et al., 2022; Chaguza et al., 2023; Markov *et al*., 2023)(Figure 1A), particularly in viral genes that code for surface proteins; (2) Multiple co-circulating viral lineages often establish within individuals with prolonged infection (Rogers *et al*., 2015; Chaguza et al., 2023; Machkovech et al., 2024) (Figure 1B); (3) Parallel viral substitutions often occur across individuals experiencing prolonged infection (Memoli *et al*., 2014; Xue et al., 2017; Wilkinson et al., 2022) (Figure 1C); (4) The extent of antigenic evolution that is observed across individuals with prolonged infection is highly variable (Rocha *et al*., 1991; McMinn et al., 1999; Xue et al., 2017; Harari *et al*., 2022) (Figure 1D).

**Figure 1.**
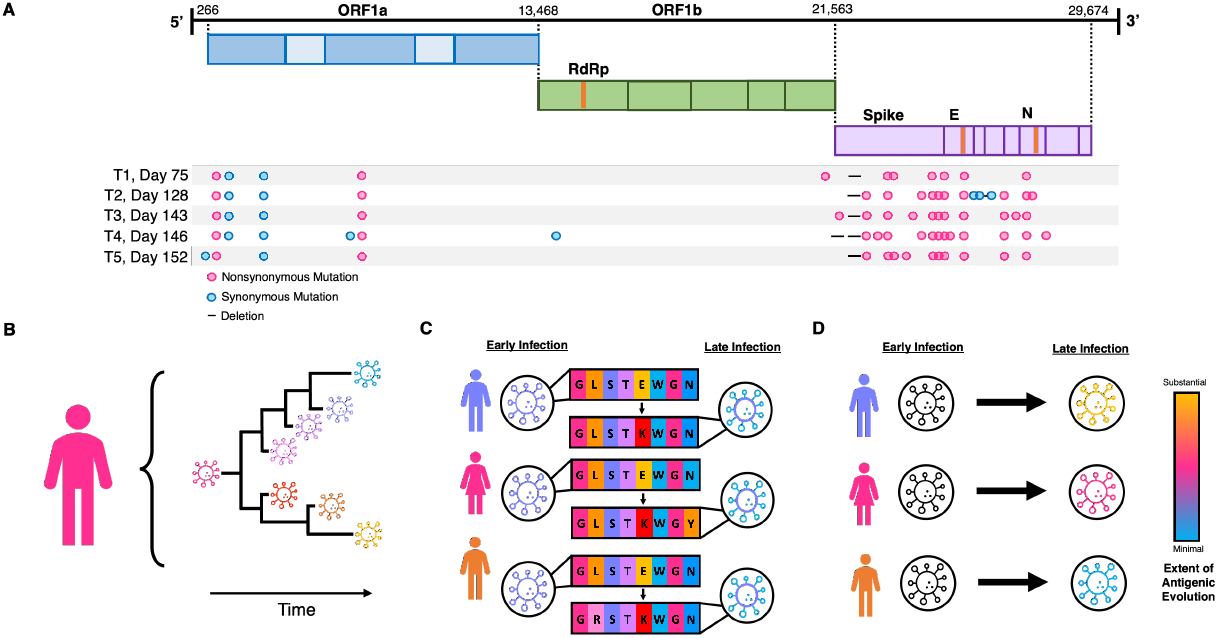
Evolutionary patterns of respiratory viruses observed in individuals with prolonged infection. (A) Nonsynonymous substitution rates generally exceed synonymous substitution rates. Here, nonsynonymous and synonymous substitutions that are observed in sequences sampled over time from an individual with a prolonged SARS-CoV-2 infection are shown. Sample days and substitutions are relative to the individual’s T0, Day 0 consensus sequence. Schematic reproduced from Choi et al. (2020). (Permission to reproduce will be obtained from NEJM upon manuscript acceptance.) (B) Multiple distinct viral lineages can evolve within individuals with prolonged infections. (C) Parallel substitutions often arise across individuals with prolonged infection. The schematic shows consensus viral sequences from early and late infection timepoints of three infected individuals. Arrows highlight the frequently observed E484K substitution in the spike gene of SARS-CoV-2. (D) Rates of antigenic evolution are variable across individuals with prolonged infection. The schematic depicts heterogeneity in the extent of antigenic evolution that occurs over time in three individuals.

While these four patterns of respiratory virus evolution in individuals with prolonged infection are well established, we still lack a comprehensive understanding of their drivers. Here, we develop a simulation model for respiratory virus evolution within individuals with prolonged infection and simulate this model under various parameterizations to help shed light on possible drivers of within-host viral evolution in these longer-term infections. Specifically, we extend a tunably rugged fitness landscape model (Aita *et al*., 2000; Neidhart et al., 2014) to consider viral evolution at sites that are either synonymous or nonsynonymous, with the latter impacting either viral replicative fitness, antigenicity, or both replicative fitness and antigenicity (sites that we call pleiotropic sites). We simulate this model to identify features of the viral fitness landscape and characteristics of the immune response that can reproduce these four evolutionary patterns that have been empirically observed in prolonged respiratory virus infections. From these simulations, we find that immune pressure is key to consistently reproducing all four viral evolutionary patterns shown in Figure 1.

## Methods

### The Fitness Landscape Model

We model the viral genome as consisting of four different types of sites: synonymous sites (S), phenotypic sites (P), antigenic sites (A), and pleiotropic sites (PA) (Figure 2A). Mutations falling on synonymous (S) sites are assumed to be silent and do not impact viral fitness. Mutations falling on phenotypic sites are nonsynonymous and impact phenotypes related to replicative fitness. Mutations falling on antigenic sites are nonsynonymous and impact only antigenicity. Mutations falling on pleiotropic sites are nonsynonymous and simultaneously impact both replicative fitness and antigenicity. The number of sites in each of these four classes is given by *L*_*S*_, *L*_*P*_, *L*_*A*_, and *L*_*PA*_, respectively, with the total number of sites in the viral genome given by *L* = *L*_*S*_ + *L*_*P*_ + *L*_*A*_ + *L*_*PA*_. Genotypes are modeled as bitstrings, with each site carrying one of two possible alleles: 0 or 1. As such, genotype space consists of 2^*L*^ genotypes. We determine the overall fitness of a given viral strain by taking into consideration its replicative fitness as well as its antigenic phenotype.

**Figure 2.**
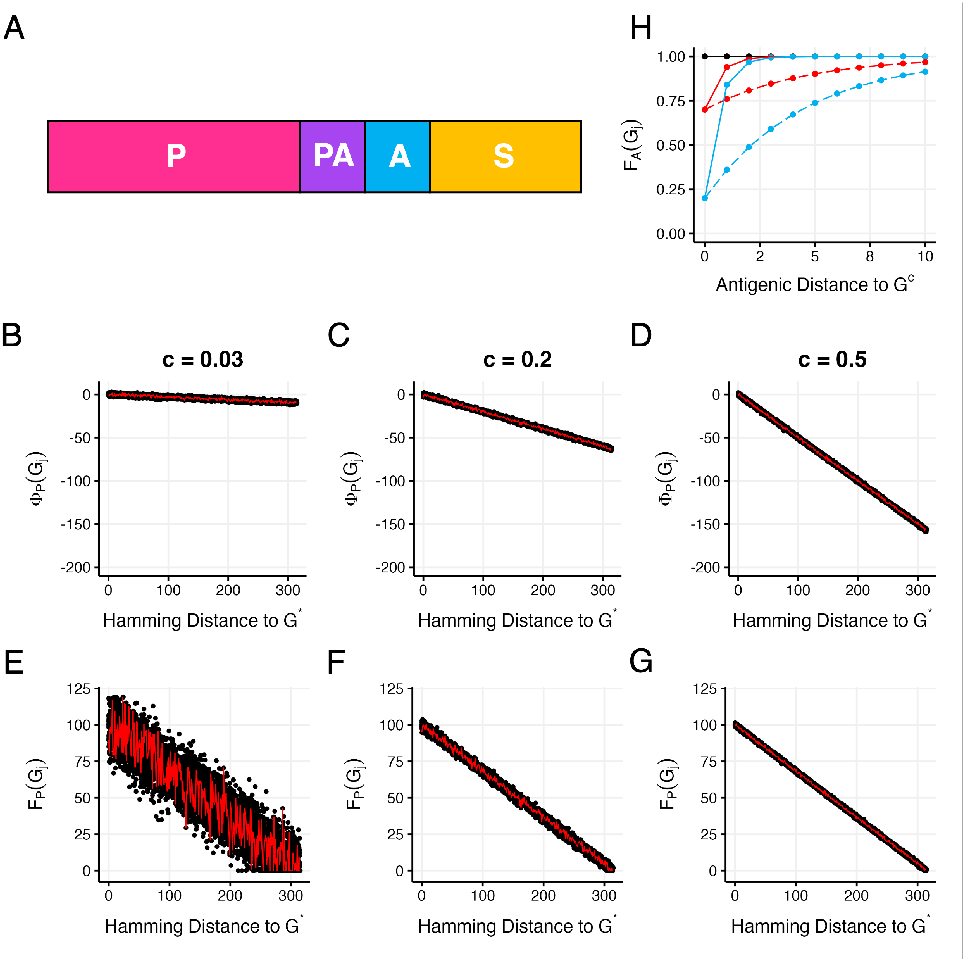
Structure of the viral genome and projections of replicative fitness and antigenic fitness. (A) Structure of the viral genome, showing the 4 different types of sites: phenotypic (*P*), pleiotropic (*PA*), antigenic (*A*), and synonymous (*S*). Site order is not important because we do not consider the process of recombination in our simulations of viral evolution. (B) Projection of the Rough Mount Fuji fitness landscape given by equation (1), parameterized with a small *c* to implement a highly rugged fitness landscape. Here, *c* = 0.03. (C) Projection of the RMF fitness landscape given by equation (1), parameterized with a *c* of 0.2 to implement a semi-rugged fitness landscape. (D) Projection of the RMF fitness landscape with a large *c* to implement a relatively smooth fitness landscape. Here, *c* = 0.5. In (B-D), *G*^*\**^ is given by the bitstring of all ones and the genome length is set to *L*_*P*_ + *L*_*P A*_ = 315. (E-G) Linearly transformed fitness landscapes, derived from panels (B-D), with parameter *k* set to 100. The red lines in panels (B-G) show one genetic pathway consisting of single point mutations from the bitstring of all ones (*G*^*\**^; Hamming distance to *G*^*\**^ = 0) to the bitstring of all zeros (Hamming distance to *G*^*\**^ = 315). H) Antigenic fitness of genotypes that are different antigenic distances away from the consensus genotype *G*^*c*^. The black line shows the antigenic fitness function parameterized with *q* = 0 (and any value of *p*). The red line shows *F*_*A*_ with *q* = 0.3 and *p* = 0.2, corresponding to immune strength being weak and immune breadth being narrow. The red dashed line shows *F*_*A*_ with *q* = 0.3 and *p* = 0.8, corresponding to immune strength being weak and immune breadth being moderate. The blue line shows *F*_*A*_ with *q* = 0.8 and *p* = 0.2, corresponding to immune strength being strong and immune breadth being narrow. The blue dashed line shows *F*_*A*_ with *q* = 0.8 and *p* = 0.8, corresponding to immune strength being strong and immune breadth being moderate.

To quantify the replicative fitness of a viral genotype, we extend an existing fitness landscape model called the Rough Mount Fuji (RMF) model (Aita *et al*., 2000; Neidhart *et al*., 2014). The original model has two parameters: the length of the genome and a parameter *c* that controls the ruggedness of the landscape. The landscape is a static landscape, with the fitness of a genotype not changing over time. We thus use the RMF landscape only for quantifying the replicative fitness component of a viral genotype. Because in our model replicative fitness depends only on the alleles present at P and PA sites, the length of the genome we use for the RMF model is given by *L*_*P*_ + *L*_*PA*_, resulting in a total of 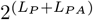 possible genotypes on the landscape. Fitness values in the RMF model are initially calculated according to the following equation (Neidhart *et al*., 2014):

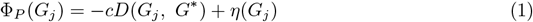

where *G*_*j*_ denotes focal genotype *j* and *G*^*\**^ denotes a reference genotype. The first term, *−cD*(*G*_*j*_, *G*^*\**^), is a product of the non-negative parameter *c* and the Hamming distance between genotype *j* and reference genotype *G*^*\**^. The Hamming distance is defined as the number of sites at which the two genotypes differ. The minimum value of *D* is 0, which occurs when *G*_*j*_ = *G*^*\**^. The maximum value of *D* is *L*_*P*_ + *L*_*PA*_, which occurs when focal genotype *G*_*j*_ differs from *G*^*\**^ at every site that contributes to replicative fitness. The second term, *η*(*G*_*j*_), is a discrete random variable that is drawn from a specified probability distribution function. Here, we use a Gaussian distribution with mean 0 and standard deviation 1 to generate our *η* random variables. This RMF model generates a tunably rugged fitness landscape model, with the random variable *η*(*G*_*j*_) contributing the ruggedness and the parameter *c* modulating the extent of the ruggedness. When *c* = 0, the fitness landscape reduces to a House of Cards landscape, where the fitness value of a genotype is uncorrelated with the fitness value of genotypes that surround it. At low *c* (high ruggedness), epistatic interactions dominate. As *c* gets larger, epistatic interactions become weaker and the fitness landscape becomes less rugged, reducing to a smooth Mount Fuji landscape as *c → ∞*, with genotype *G*^*\**^ being the genotype with highest fitness. Figures 2B-D project the RMF landscape for three values of *c*: *c* = 0.03, *c* = 0.2, and *c* = 0.5.

Under this model, the range of values that the fitness landscape spans differs depending on the value of *c* chosen. When *c* is larger, the range of fitness values is larger (Figures 2B-D). To be able to compare patterns of viral evolution across fitness landscapes of different ruggedness, we therefore modified the above RMF model such that the overall distribution of fitness values would be similar across *c* values. We did this by linearly transforming the Φ_*P*_ fitness values so that that they fall between 0 and *k*, where *k* is a non-negative parameter, similar to the approach taken in Greene and Crona (2014). Because some of the genotypes that are distant from the reference genotype may have fitness values that fall below 0 following linear transformation, we set the fitness values of these genotypes to 0. We denote the linearly-transformed replicative fitness value of genotype *G*_*j*_ as *F*_*P*_ (*G*_*j*_). Figures 2E-G project the linearly-transformed fitness landscape for the RMF landscapes shown in Figures 2B-D, respectively, with *k* set to 100. These linearly-transformed fitness landscape projections demonstrate that a lower value of the parameter *c* results in a more rugged fitness landscape, with many local fitness peaks present across the landscape.

We allow antigenic changes (from mutations occurring at either antigenic (*A*) sites or pleiotropic (*PA*) sites) to impact overall viral fitness. Specifically, we assume that the overall fitness of a viral genotype *G*_*j*_ depends on both viral replicative fitness and its antigenic fitness:

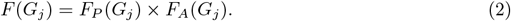

Here, antigenic fitness *F*_*A*_(*G*_*j*_) is a value between 0 and 1. When *F*_*A*_(*G*_*j*_) is closer to 0, there is substantial immune pressure against genotype *G*_*j*_ that results in its overall fitness being close to 0. As *F*_*A*_(*G*_*j*_) approaches 1, immune pressure against genotype *G*_*j*_ is weaker and overall fitness is determined solely by replicative fitness. By modeling antigenic fitness in this manner, we are assuming that it modulates viral fitness by introducing a fitness cost. We model antigenic fitness using the function:

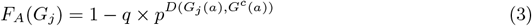

where *G*^*c*^ denotes the consensus genotype circulating in the viral population and *D*(*G*_*j*_(*a*), *G*^*c*^(*a*)) denotes the Hamming distance between genotype *G*_*j*_ and consensus genotype *G*^*c*^ at exclusively antigenic and pleiotropic sites (notationally referred to here as *a* sites). This function contains 2 parameters. Taking on values between 0 and 1, parameter *q* can be thought of as a parameter that modifies the *strength* of immune pressure. When *q* = 0, the antigenic fitness of the consensus genotype (and all genotypes) is 1, reflecting an absence of immune pressure. As *q* approaches 1, the antigenic fitness of the consensus genotype approaches 0, indicating that immune pressure is strong and considerably impacts overall viral fitness. Parameter *p* can be thought of as a parameter that modifies the *breadth* of immunity. Also taking on values between 0 and 1, when *p* is closer to 0, the breadth of the immunity is narrow, with genotypes that are only a single antigenic mutation away from the consensus genotype experiencing only very weak immune pressure (their *F*_*A*_ values being close to 1). As *p* approaches 1, immune protection becomes broader, with genotypes that are several antigenic mutations away from the consensus genotype still experiencing similar immune pressure to that of the consensus genotype. Figure 2H shows the antigenic fitness values of viral genotypes that are different antigenic distances away from the consensus genotype under different parameterizations of *q* and *p*.

A mutation at a synonymous site that generates viral genotype *G*_*j*_ does not impact either *F*_*P*_ (*G*_*j*_) or *F*_*A*_(*G*_*j*_), such that the overall fitness of the mutant is the same as its parent. A mutation at a phenotypic *P* site impacts only *F*_*P*_ (*G*_*j*_). A mutation at an antigenic *A* site impacts only *F*_*A*_(*G*_*j*_). A mutation at a pleiotropic *PA* site impacts both *F*_*P*_ (*G*_*j*_) and *F*_*A*_(*G*_*j*_). Finally, it is important to note that, by the formulation of equation (3), we are assuming that immune escape is fleeting in that viral fitness with respect to antigenicity only depends on how different a genotype is from the consensus genotype at the present time and not the history of viral genotypes that have circulated over the course of infection.

### Simulating within-host viral evolution

We simulate within-host viral evolution under the assumption of a fixed viral population size *N* using Gillespie’s *τ* -leap algorithm (Gillespie, 2001). Initially, at the time of infection (*t* = 0), each viral particle is set to the same infecting genotype. We adopt this assumption to reflect the low levels of viral diversity at the start of infection that result from a tight transmission bottleneck (McCrone *et al*., 2018; Martin and Koelle, 2021; Lythgoe *et al*., 2021; Shi et al., 2024). We then update the evolving viral population from one time point to the next by determining which viral particles will die and which viral particles will reproduce over the next Δ*t* time increment. To determine which viral particles will die, we first draw a random number *n* from a Poisson distribution with mean *dN* Δ*t*, where *d* is the per capita viral death rate. We then choose *n* viral particles at random and without replacement to be removed from the viral population. To determine which viral particles will reproduce, we draw *n* viral particles (with replacement) in proportion to their overall fitness values *F*. Progeny viruses inherit the genotype of their respective parent, plus any additional mutations that occur during their ‘birth’. The number of additional mutations that occur during the birth of a given viral particle is drawn from a Poisson distribution with mean *µL*, where *µ* is the per site per infection cycle mutation rate and *L* is the total number of sites in the viral genome. The sites at which these mutations occur are selected at random from the viral genome. Once viral particle deaths and births have been updated, time is incremented from *t* to *t* + Δ*t*.

When simulating viral evolution using a small number of sites, it is possible to *a priori* calculate the replicative fitness values of each genotype prior to simulating viral evolution on the landscape. However, even with only 100 sites that impact replicative fitness, genotype space becomes too large to adopt this approach. When simulating viral evolution, we therefore dynamically allocate *F*_*P*_ fitness values as strains are accessed through mutation over the course of a prolonged infection. Adopting this approach allows us to calculate and store a considerably smaller portion of viral genotype space.

### Model Parameterization

We parameterize the evolutionary model using a total of *L* = 400 sites. We let 315 of these sites be nonsynonymous and the remaining 85 sites be synonymous (*S*). This ratio of 315:85 mirrors the approximate ratio of 3.7 in nonsynonymous to synonymous mutations observed for respiratory viruses such as coronaviruses and influenza viruses. Of the 315 nonsynonymous sites, we consider in our simulations different scenarios for these sites being classified as phenotypic (*P*) or pleiotropic (*PA*) sites. We do not consider any sites to be only antigenic (*A*) because existing studies of respiratory viruses that have evaluated the replicative fitness of mutations along with their antigenic impact indicate that few mutations impact only antigenicity (Greaney *et al*., 2022; Bloom and Neher, 2023; Dadonaite et al., 2024). Although we set *L*_*A*_ to zero in all of our simulations, our model still implements these sites to allow for generality. In all of our simulations, we set the parameter *k* to 100, such that replicative fitness spans values from 0 to 100 (or a little higher for the must rugged landscapes considered; see Figure 2E). Unless otherwise noted, we further set infecting genotypes to be approximately 50% adapted to the host by setting the reference genotype *G*^*\**^ to all ones and letting infecting genotypesbe bitstrings that contain 50% ones and 50% zeros at randomly chosen sites across their genomes.

We set the mutation rate to *µ* = 2.5 *×* 10^*−*5^ mutations per site per infection cycle. This mutation rate lies between the estimated mutation rate of approximately 10^*−*6^ mutations per site per cycle for coronaviruses (Bar-On *et al*., 2020; Amicone et al., 2022) and the estimated mutation rate of approximately 10^*−*4^ mutations per site per cycle for influenza A viruses (Pauly *et al*., 2017). At this mutation rate and with a genome length of *L* = 400, two or more mutations are expected to occur in less than 0.005% of replications. As such, in our simulations, we only allow zero mutations (with probability *e*^*−µL*^ *≈* 0.99) or one mutation (with probability 1 *− e*^*−µL*^ *≈* 0.01) to occur during the process of viral replication. We set the generation time to 6 hours, corresponding to a death rate of *d* = 4 per day. We chose this generation time based on the 6-8 hour generation time estimated for influenza viruses (Einav *et al*., 2020) and the 6-9 hour generation time estimated for SARS-CoV-1 (Schneider *et al*., 2012; Bar-On et al., 2020). In all of our simulations, we calculate overall fitness values for genotypes using equation (2) and set the time step to Δ*t* = 1 hour.

## Results

### Viral population size modulates the strength of selection and genetic drift

We first simulated the model to confirm that it recovers the well-established evolutionary pattern that genetic drift dominates when (effective) population sizes are small and that selection acts more efficiently when these population sizes are larger. To this end, we simulated viral evolution on a relatively smooth replicative fitness landscape (*c* = 0.5) for two different viral population sizes: *N* = 40 and *N* = 5000. The viral genome in these simulations consisted of 85 synonymous (*S*) sites and 315 phenotypic (*P*) sites. Figure S1 shows the evolutionary dynamics of simulated viral populations over the time course of 3 years. When viral population sizes are small (*N* = 40), mean fitness of these populations does not consistently increase (Figure S1A), indicating a lack of viral adaptation. In contrast, when viral population sizes are larger (*N* = 5000), mean population fitness increases (Figure S1B). In small populations, genetic divergence accrues at a rate that is similar to that expected under neutral evolution (Figure S1C), whereas in large populations, genetic divergence accrues at a rate that exceeds that expected under neutral evolution (Figure S1D), again, indicating the occurrence of viral adaptation in these large populations. Finally, as one would expect, the average pairwise genetic diversity is lower in the small viral populations (Figure S1E) than in the large viral populations (Figure S1F). More interestingly, the small populations show patterns of genetic diversity that are consistent with expected levels of genetic diversity under neutral evolution (Figure S1E), whereas the large populations show patterns of genetic diversity that are substantially lower than those expected under neutral evolution (Figure S1F). This is consistent with positive selection acting to reduce genetic diversity in the large-*N* simulations. Together, the results shown in Figure S1 indicate that viral population sizes modulate the strength of selection and genetic drift, with genetic drift dominating in small viral populations and selection dominating in large viral populations, as expected from population genetic theory.

### Observed excess of nonsynonymous substitutions is incompatible with viral evolution on static fitness landscapes

We next used our model to explore the impact that fitness landscape ruggedness has on patterns of within-host viral evolution and adaptation, specifically focusing on what types of fitness landscapes could consistently reproduce the empirically observed pattern that nonsynonymous substitution rates generally exceed synonymous substitution rates in prolonged viral infections (Figure 1A). To this end, we considered four fitness landscapes across a range of ruggedness from *c* = 0.03 (highly rugged) to *c* = 0.5 (smooth). In each case, we set the the viral population size to *N* = 5000 based on findings that viral effective population sizes in prolonged infections are thought to be large (Xue *et al*., 2017; Lumby *et al*., 2020), while maintaining computational tractability. We further considered only synonymous sites and sites impacting replicative fitness (*L*_*S*_ = 85, *L*_*P*_ = 315, *L*_*A*_ = 0, *L*_*PA*_ = 0) and simulated viral evolution on each of the four fitness landscapes for one-year periods. Six independent replicates were simulated for each fitness landscape to be able to assess general trends.

On a highly rugged landscape (*c* = 0.03), viral populations rapidly adapted in the first 2 months following infection, from a mean fitness value of approximately 50 to a mean fitness value of approximately 80 (Figure 3A). Following this initial bout of adaptation, further increases in fitness were less pronounced and only occurred sporadically. Divergence from the infecting genotype increased rapidly during the initial period of adaptation but then tended to slow down (Figure 3E), with divergence considerably lower than expected under neutral evolution during later time points. These results are consistent with initial movements of the viral populations to local higher-fitness peaks in the landscape, followed by the populations getting ‘trapped’ in these local peaks. Indeed, when we plot changes in these populations’ consensus sequences over time, we see that a small number of nonsynonymous substitutions occurred shortly following infection, but additional nonsynonymous substitutions were rare thereafter (Figure 3I). In contrast, synonymous substitutions continue to accumulate over the year of simulation (Figure 3I). Figure 3I further indicates that nonsynonymous viral substitution rates would be expected to be lower than synonymous ones in viral populations that have evolved on this rugged fitness landscape when calculated 6-12 months following infection. We next simulated viral evolution on a less rugged fitness landscape (*c* = 0.1). Mean fitness of the simulated viral populations also rapidly increased within the first 2 months of infection, but fitness levels only reached values of approximately 60 (Figure 3B), rather than the fitness values of 80 that were observed on the more rugged fitness landscape of *c* = 0.03. This indicates that viral populations still got trapped in local fitness peaks on this less rugged landscape, despite the landscape being smoother. The remainder of the evolutionary patterns on the *c* = 0.1 landscape are highly similar to those observed on the *c* = 0.03 landscape: divergence tended to level off (Figure 3F) and nonsynonymous substitutions occurred only early on in infection and then purifying selection dominated (Figure 3J), resulting in fewer nonsynonymous substitutions than synonymous substitutions by 6-12 months post-infection. Patterns of viral evolution on an even smoother fitness landscape (*c* = 0.2) look similar to those on the *c* = 0.1 landscape, with the only appreciable difference being that the initial bouts of viral adaptation that occurred during the first 2 months of infection reached even lower fitness plateaus (55-60, rather than approximately 60). Finally, on the smoothest fitness landscape considered (*c* = 0.5), fitness levels continued to increase over the simulated year-long infections (Figure 3D). These continuous increases, however, only increased mean fitness slightly. Unlike the populations that evolved on the more rugged landscapes, divergence in the populations evolving on the smooth *c* = 0.5 landscape continued to increase, consistent with neutral patterns of divergence (Figure 3H). Nonsynonymous substitutions also continued to accrue, but only at rates similar to those of synonymous substitutions (Figure 3L). This is because the fitness impacts of phenotype-impacting mutations on these smooth landscapes were small, approaching nearly-neutral. Nevertheless, selection was still able to act (albeit slowly) on the viral populations evolving on these smooth landscapes, as indicated by the consistent increase in mean fitness in these populations (Figure 3D).

**Figure 3.**
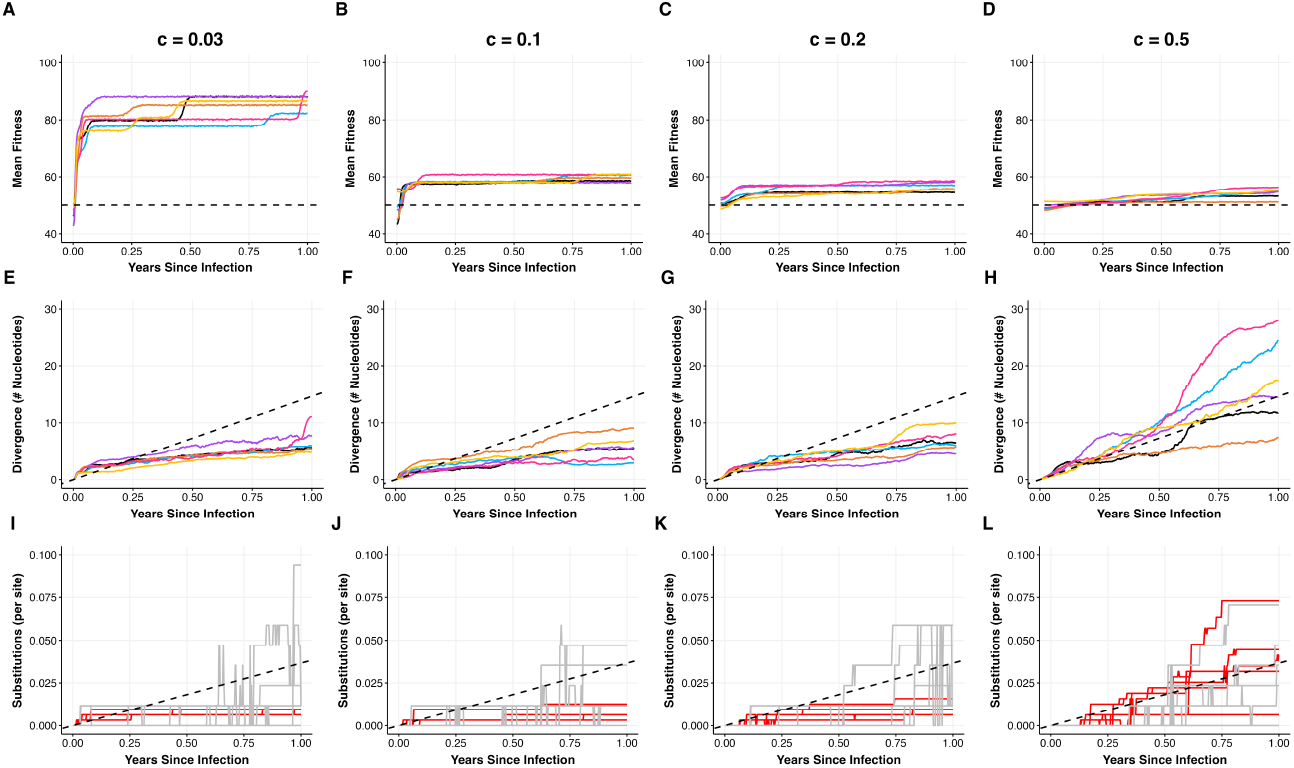
Patterns of viral adaptation observed across fitness landscapes of variable ruggedness. Columns correspond to simulations of viral populations on fitness landscapes of increasing smoothness, with the *c* = 0.03 landscape implementing a highly rugged landscape (first column) and the *c* = 0.5 landscape (last column) implementing a relatively smooth landscape. (A-D) Mean population fitness over the course of infection for six independent simulations. The dashed black line at 50 indicates the expected fitness of the infecting genotype. (E-H) Mean divergence from the infecting genotype for the same six populations. The dashed black line shows expected divergence under neutral evolution. (I-L) Number of nonsynonymous and synonymous substitutions in the six populations over time. For a given simulation, substitutions are calculated between the consensus sequence at a given time point and the infecting genotype. Red lines show nonsynonymous substitutions. Grey lines show synonymous substitutions. Substitutions are normalized by the number of nonsynonymous and synonymous sites, respectively, yielding a per-site number of substitutions. Dashed black line shows the expected number of substitutions under neutral evolution. All simulations used a viral genome of length *L* = 400, with *L*_*S*_ = 85 and *L*_*P*_ = 315. Each infecting genotype had a Hamming distance of 200 from the reference genotype of all ones. Other parameters were: *N* = 5000, *µ* = 2.5 *×* 10^*−*5^ mutations per site per infection cycle, *k* = 100, *d* = 4 infection cycles per day.

Together, these simulations indicate that simulations of viral evolution occurring on fitness landscapes of variable ruggedness do not recapitulate patterns of nonsynonymous substitutions being in excess of synonymous substitutions.

### Viral populations evolving on static fitness landscapes do not diversify into distinct lineages during their adaptation

While the results shown in Figure 3 indicate that viral adaptation is expected to occur across fitness landscapes that differ in their ruggedness, they did not yield information on the extent to which the viral populations diversified throughout their adaptation. It is conceivable that the viral populations remained largely monomorphic, in each simulation traversing to a single local fitness peak. Alternatively, viral populations could have diversified, leading to the occupation of many different local fitness peaks. To determine which of these possibilities occurred, we serially sampled the simulated viral populations on a monthly basis and inferred neighbor-joining trees from the sampled viruses (Figure 4). Across all four fitness landscapes considered, these trees show no evidence of diversification into distinct lineages, despite some viral turnover being apparent in the phylogenies. As such, none of these simulations reproduced the empirically observed pattern of viral lineage diversification commonly observed in prolonged infections (Figure 1B). These phylogenies further indicate that viral populations within single individuals ultimately got trapped in single fitness peaks, rather than in multiple fitness peaks.

**Figure 4.**
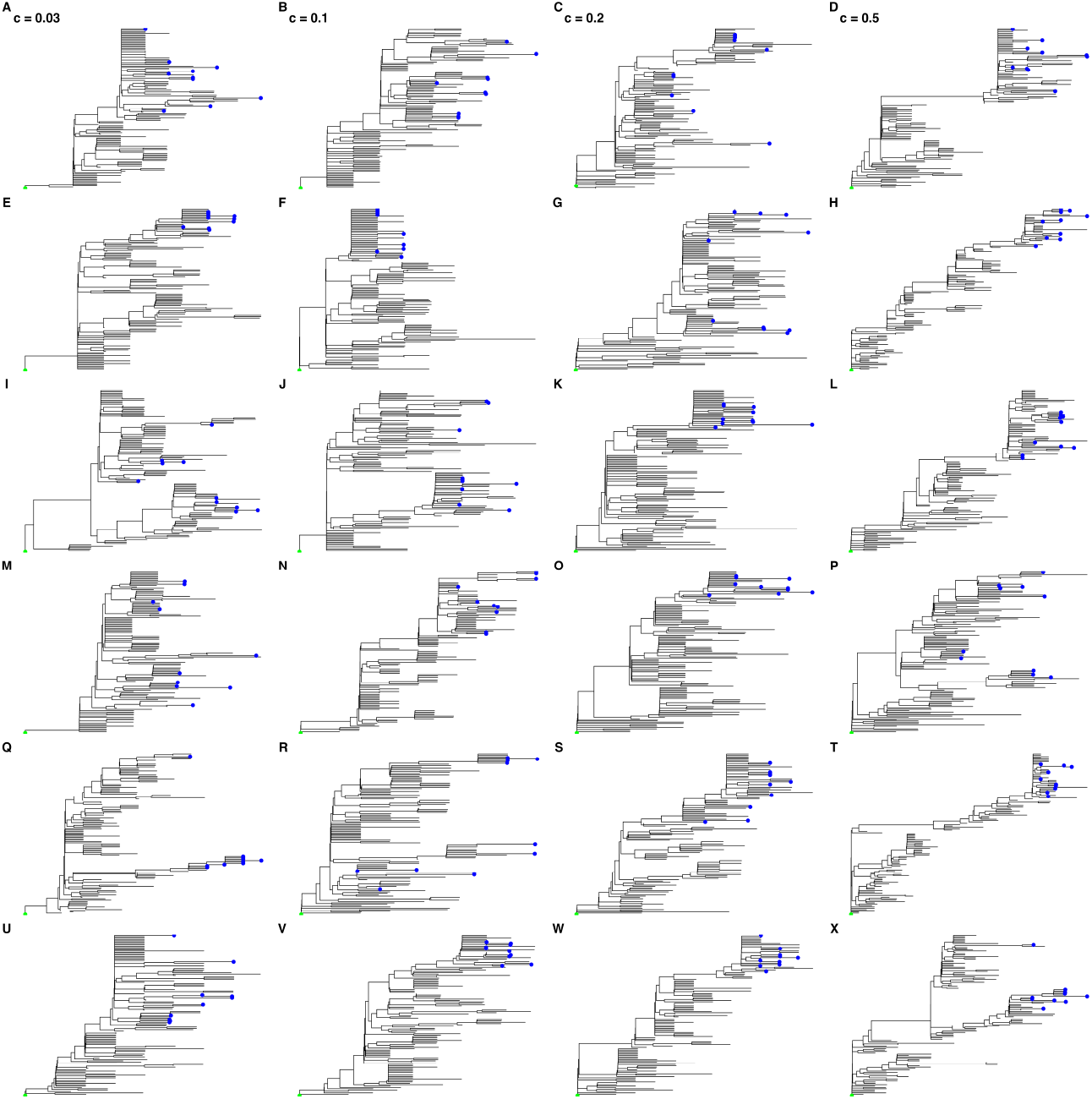
Phylogenies inferred from simulated viral populations evolving on fitness landscapes of variable ruggedness. Columns correspond to the fitness landscapes in Figure 3, ranging from a highly rugged fitness landscape (*c* = 0.03, first column) to a smooth fitness landscape (*c* = 0.5, fourth column). Rows correspond to the six independent simulations shown in Figure 3. For each simulation, a neighbor-joining tree was inferred using a sequence dataset that contained the infecting genotype as well as 10 sequences that were sampled monthly over the year-long simulation. Sequences were randomly sampled from the viral population. Neighbor-joining trees were generated using the R packages Ape (Paradis and Schliep, 2019) and treeio (Wang *et al*., 2020) and plotted using ggtree (Yu *et al*., 2017). The infecting genotype is labeled in green and viruses sampled at the last time point are labeled in blue. All trees were rooted on the infecting genotype.

### Parallel substitutions do not occur readily on static fitness land-scapes

We now address through simulation the question of whether fitness landscapes of various ruggedness can consistently reproduce the pattern of parallel substitutions that is frequently observed across individuals with prolonged infections (Figure 1C). We again considered viral evolution on the four different fitness landscapes we considered in Figure 3, and simulated viral evolution for 6 months. For each landscape, we started each of the six simulations off with the same infecting genotype that was approximately 50% adapted to the host and used the same fitness landscape for each of the simulations. As such, genotypes that had been discovered in a previous simulation and had their fitness values already dynamically allocated retained their fitness values across simulations.

On a very rugged landscape (*c* = 0.03), the mean fitness of all six simulated viral populations increased, largely in the 2 months following infection (Figure 5A). These dynamics are, as expected, consistent with the dynamics of mean fitness that were observed across different fitness landscapes parameterized with *c* = 0.03 (Figure 3A). Interestingly, the fitness levels at which the populations plateaued differed across the six simulations, despite the same underlying fitness landscape and the same infecting genotype. This indicates that the different viral populations may have found different nearby local fitness peaks. To explore this possibility, we identified, for each population, the set of sites that contained a nonsynonymous high-frequency allele that differed from the infecting genotype at 6 months post-infection, defining high-frequency as exceeding 20%. For each pair of individuals, we then determined the number of sites that were shared across their sets. On the *c* = 0.03 landscape, we found that very few high-frequency mutations (if any) were shared between individuals (Figure 5E). To understand these results, we can remember that highly rugged landscapes come close to a House of Cards landscape, where the fitness effect of a mutation depends almost entirely on its genetic context (that is, epistatic interactions dominate fitness effects). As such, one would expect the first nonsynonymous substitution to impact the fitness effects of all other possible nonsynonymous substitutions. Parallel substitutions would therefore only likely be observed if the first substitution was the same one across individuals. Once different substitutions occurred, the viral populations across the different individuals would be expected to be on different evolutionary paths due to the extreme ruggedness of the fitness landscape.

**Figure 5.**
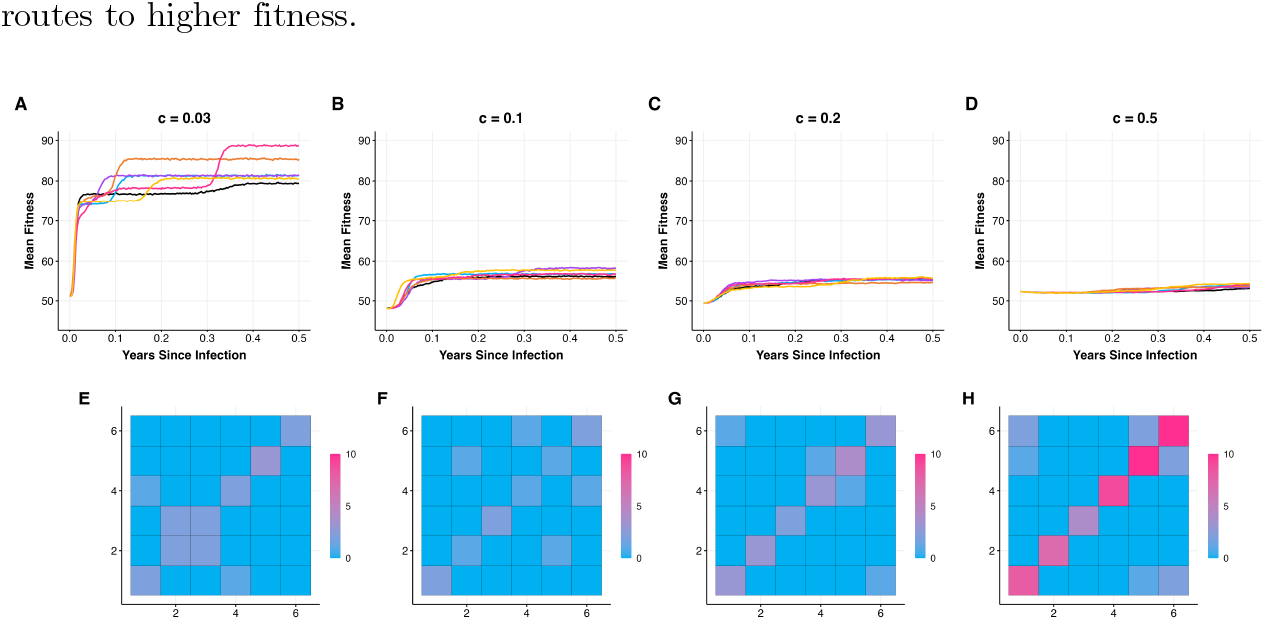
Parallel mutations do not readily occur on static fitness landscapes. (A-D) Changes in mean population fitness of six viral populations evolving on fitness landscapes of various ruggedness: (A) *c* = 0.03; (B) *c* = 0.1; (C) *c* = 0.2; (D) *c* = 0.5. On each landscape, the six viral populations each started with the same infecting genotype and evolved on the same landscape. (E-H) The number of shared mutations across pairs of individuals. Only mutations at nonsynonymous sites that exceeded frequencies of 20% at time *t* = 0.5 years were considered in this calculation. Cells along the diagonal show the number of high-frequency nonsynonymous mutations in each individual at 0.5 years post infection. Model parameters are: *N* = 5000, *µ* = 2.5 *×* 10^*−*5^ mutations per site per infection cycle, *k* = 100, *d* = 4 replications per day, *L*_*S*_ = 85 and *L*_*P*_ = 315.

On a semi-rugged landscape parameterized with *c* = 0.1, we see that fitness increases to similar levels across the individual infections (Figure 5B). However, these fitness increases appear to largely stem from different nonsynonymous substitutions, given patterns of shared nonsynonymous high-frequency mutations (Figures 5F). Viral evolution on even smoother fitness landscapes again result in only rare shared nonsynonymous variation (Figures 5G,H). These results make sense in that, on smoother landscapes, beneficial mutations all have similar fitness effects, such that there are many different routes to higher fitness.

Together, our results shown in Figures 3-5 indicate that adaptation is readily observed in simulated viral populations evolving on a range of different fitness landscapes. However, on rugged landscapes, adaptation occurs rapidly and then evolution is dominated by purifying selection, such that nonsynonymous substitution rates do not exceed synonymous ones in the longer-term. On smooth landscapes, adaptation continues to occur, but fitness differentials are so small that nonsynonymous substitution rates are similar to synonymous ones in the longer-term. As such, viral evolution on these static fitness landscapes cannot consistently reproduce the observed excess of nonsynonymous substitutions observed in prolonged infections. Our simulations on these static landscapes further indicate that co-circulating viral lineages do not readily evolve and that parallel mutations are not readily observed on any of the considered fitness landscapes. We therefore next consider the role that mutations that impact antigenicity may play in the reproduction of these observed patterns. Specifically, we consider the impact of pleiotropic sites on these patterns, where mutations at these sites impact both replicative fitness and antigenicity.

### Interindividual variation in rates of antigenic evolution can be explained by differences in the strength and breadth of immune pressure

To assess the impact that pleiotropic sites have on patterns of within-host viral evolution in prolonged infections, we assume a semi-rugged fitness landscape of *c* = 0.2, again with 315 nonsynonymous sites and *L*_*S*_ = 85 synonymous sites. We further assume that a subset of the nonsynonymous sites impact only replicative fitness (*L*_*P*_ = 267) while the remainder of the nonsynonymous sites impact both replicative fitness and antigenicity (*L*_*PA*_ = 48). We consider different strengths of immune pressure as well as different breadths of the immune response. Our first simulations consider variation in the strength of immune pressure, modified by varying the value of parameter *q* in equation (3). Figure 6 shows simulations at four different values of *q*, ranging from *q* = 0.0 (no immune pressure) to *q* = 0.9 (very strong immune pressure). For all simulations we kept immune breadth constant at a moderate level of *p* = 0.8. The strength of immune pressure (*F*_*A*_) adopted under these scenarios is shown graphically in Figure S2.

**Figure 6.**
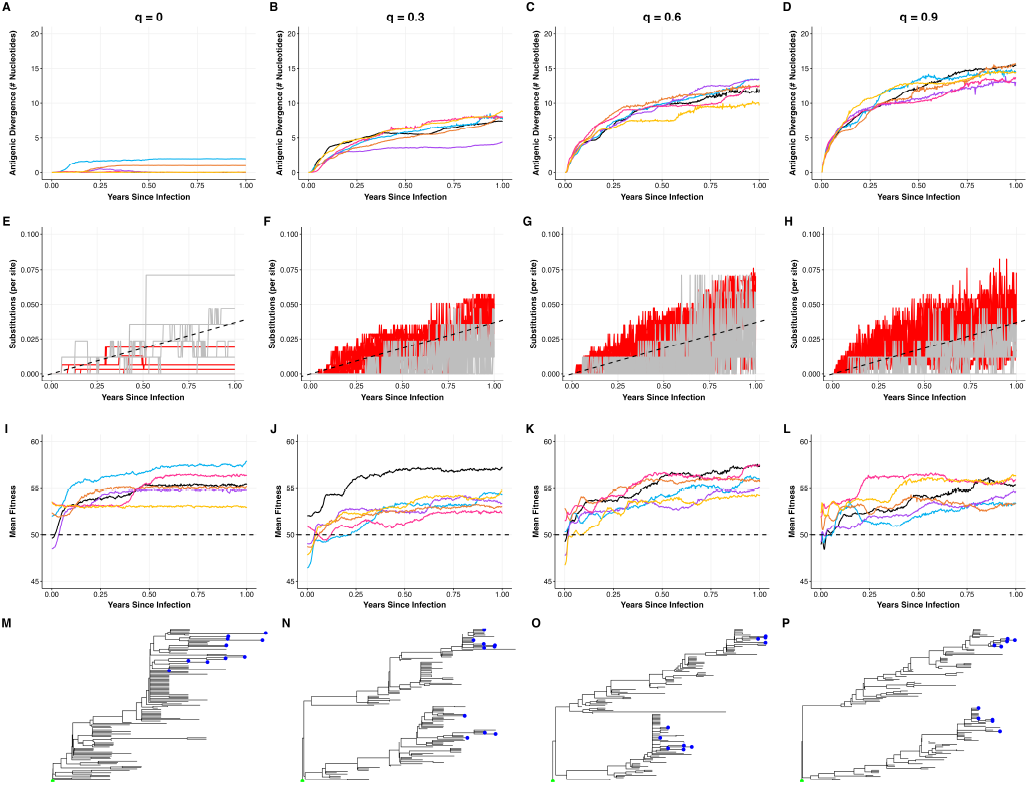
The strength of the immune response impacts patterns of within-host viral evolution in prolonged infections. Columns correspond to varying strengths of the immune response: no immune response (*q* = 0.0), low strength (*q* = 0.3), medium strength (*q* = 0.6), and high strength (*q* = 0.9). All simulations set the breadth of the immune response to *p* = 0.8. (A-D) Extent of antigenic evolution over the course of infection for six simulations. Antigenic evolution was calculated as divergence between the consensus genotype at a given time point and the infecting genotype, at the subset of nonsynonymous sites that impacted antigenicity. (E-H) Number of nonsynonymous (red) and synonymous (grey) substitutions per site over the course of infection. Dashed black line shows the expected number of substitutions under neutral evolution. (I-L) Mean viral replicative fitness over the course of infection. The horizontal dashed line shows the expected fitness of the infecting genotype. (M-P) Neighbor-joining trees, generated using 10 randomly sampled genotypes every 30 days. NJ trees are shown for the the simulation shown in black in the above panels. All trees were rooted on the infecting genotype. Simulations were performed using a viral genome of length *L* = 400, with *L*_*S*_ = 85, *L*_*P*_ = 267, *L*_*PA*_ = 48, and *L*_*A*_ = 0. Other parameters are: *N* = 5000, *µ* = 2.5 *×* 10^*−*5^ mutations per site per infection cycle, *k* = 100, *c* = 0.2, and *d* = 4 infection cycles per day.

We first performed six simulations in the absence of immune pressure (*q* = 0.0). Under this parameterization, antigenic divergence (via mutations occurring at pleiotropic sites) remained low (Figure 6A), as one might expect given the lack of immune pressure acting on these simulated viral populations. In the presence of immune pressure (*q >* 0), antigenic divergence increased over the simulated infections, with higher rates of antigenic divergence observed with stronger immune pressure (Figure 6B-D). While our results show that there is some variation in the amount of antigenic evolution that occurs within simulations that are parameterized with the same strength of immune pressure, there is considerably more variation in the amount of antigenic evolution observed across simulations that differ in the strength of immune pressure. Observed interindividual heterogeneity in the rate of antigenic evolution could thus be explained by variation across individuals in the strength of their immune response, with individuals that exert higher immune pressure giving rise to viral populations that have undergone more antigenic evolution. Of note, these simulations do not consider the impact of immune pressure on viral population sizes; with higher immune pressure, it could be the case that viral population sizes are reduced, which would decrease the efficiency of selection and therefore may reduce the rate of within-host antigenic evolution, as has been suggested previously (Grenfell *et al*., 2004).

Figure 7 considers the impact that the breadth of the immune response has on the rate of antigenic evolution. We parameterize the simulations for this figure using different values of *p* (see equation (3)), ranging from *p* = 0.0 (very narrow immune breadth) to *p* = 0.95 (broad immune breadth). In these simulations, we keep the strength of immune pressure the same across simulations at a moderate value of *q* = 0.5. The extent of immune pressure (*F*_*A*_) adopted under these scenarios is shown graphically in Figure S3. In simulations with narrow immune breadth (*p* = 0.0), antigenic evolution occurred, but only at a very slow rate (Figure 7A). As immune breadth increased (higher *p*), the rate of antigenic evolution increased (Figures 7B,7C). These results make sense in that viral genotypes that are more than 1 antigenic mutation away from the consensus genotype have incrementally higher antigenic fitness (*F*_*A*_) when immune breadth is intermediate, thereby facilitating antigenic divergence. As immune breadth increases further (towards *p* = 1.0), the fitness differential conferred by antigenicity-impacting mutations is reduced. We therefore see a reduced rate of antigenic evolution at very broad immune breadths (Figure 7D). Our results indicate that observed interindividual heterogeneity in the rate of antigenic evolution could be explained by variation across individuals in the breadth of their immune response, with individuals that have an intermediate immune breadth expected to give rise to viral populations that have undergone the most antigenic evolution. Again, these conclusions are based on results that assume that viral population sizes are not impacted by the breadth of the immune response.

**Figure 7.**
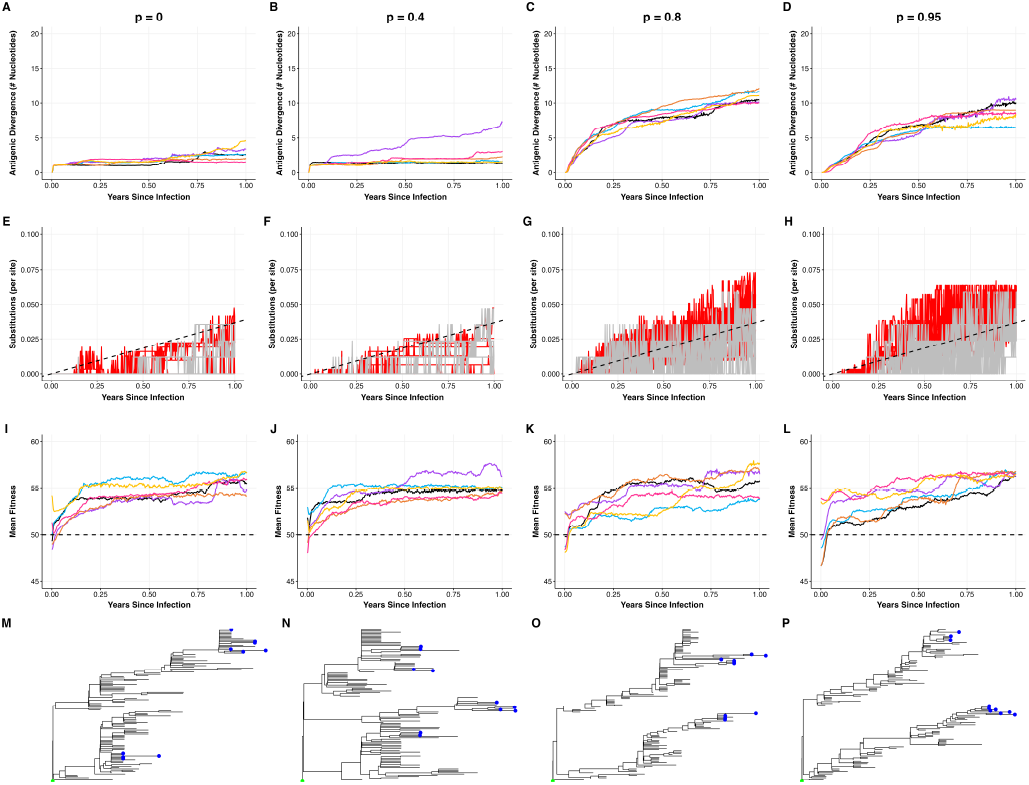
The breadth of the immune response impacts patterns of within-host viral evolution in prolonged infections. Columns correspond to varying breadths of the immune response: very narrow breadth (*p* = 0.0), low breadth (*p* = 0.4), medium breadth (*p* = 0.8), and broad breadth (*p* = 0.95). All simulations assumed a moderate strength of immune pressure (*q* = 0.5). (A-D) Extent of antigenic evolution over the course of infection for six simulations. (E-H) Number of nonsynonymous (red) and synonymous (grey) substitutions per site over the course of infection. Dashed black line shows the expected number of substitutions under neutral evolution. (I-L) Mean viral replicative fitness over the course of infection. The horizontal dashed line shows the expected fitness of the infecting genotype. (M-P) Neighbor-joining trees, generated using 10 randomly sampled genotypes every 30 days. NJ trees are shown for the simulation shown in black in the above panels. All trees were rooted on the infecting genotype. Simulations were performed using a viral genome of length *L* = 400, with *L*_*S*_ = 85, *L*_*P*_ = 267, *L*_*PA*_ = 48, and *L*_*A*_ = 0. Other parameters are: *N* = 5000, *µ* = 2.5 *×* 10^*−*5^ mutations per site per infection cycle, *k* = 100, *c* = 0.2, and *d* = 4 infection cycles per day.

In sum, our results shown in Figures 6A-D and Figures 7A-D indicate that differences between individuals in the strength and/or breadth of their immune response can reproduce heterogeneities in the extent of antigenic evolution that are observed across individuals experiencing prolonged infections (Figure 1D).

### Pleiotropic sites increase nonsynonymous substitution rates

We now revisit the patterns of viral evolution shown in Figures 1A-C to assess whether simulations with pleiotropic sites can consistently reproduce them. Figures 6E-H indicate that when the strength of the immune response is greater (larger *q*), nonsynonymous substitution rates increase and exceed synonymous substitution rates. This result makes sense in that mutations at pleiotropic sites gain a selective advantage in the presence of immune pressure, with the selective advantage being larger with stronger immune pressure. Figures 7E-H indicate that when immune breadth is at an intermediate level (moderate *p*), nonsynonymous substitution rates are higher than they are at either narrow breadth or broad breadth, and again exceed synonymous substitution rates. This result similarly makes sense in that mutations at pleiotropic sites experience the largest selective advantage when immune breadth is at an intermediate level.

While immune escape, occurring through mutations at pleiotropic sites, consistently increases nonsynonymous substitution rates, its impact on viral replicative fitness is varied. When infecting genotypes are of average fitness (approximately 50% adapted), mean replicative fitness increases similarly across simulations without immune pressure (Figure 6I) as with immune pressure of various strengths and breadths (Figures 6J, 6K, 6L and Figures 7I, 7J, 7K, 7L). Under these scenarios, therefore, immune escape neither impedes nor facilitates viral phenotypic adaptation.

The presence of host immune pressure, however, does impact viral fitness unrelated to antigenicity when the infecting genotype is either poorly adapted to the host or when it is well-adapted to the host (Figures S4, S5). When the infecting genotype is poorly adapted to the host, immune pressure tends to facilitate viral phenotypic adaptation, with mean fitness levels increasing to higher levels than in the absence of immune pressure (Figure S4). In contrast, when the infecting genotype is well adapted to the host, immune pressure tends to impede viral phenotypic adaptation, with mean fitness levels evolving to be lower than in the absence of immune pressure (Figure S5). These results make sense in terms of immune pressure driving antigenic evolution. When antigenic evolution occurs at pleiotropic sites, then these evolutionary dynamics increase mean replicative fitness if the replicative fitness effect of these mutations are on average positive, which they are if the infecting genotype is poorly adapted. In contrast, these evolutionary dynamics decrease mean replicative fitness if the replicative fitness effect of these mutations are on average negative, which they are in the infecting genotype is well adapted. As such, we would not expect immune pressure to either always facilitate or impede viral adaptation. The net impact of immune pressure on viral adaptation unrelated to antigenicity is expected to depend on the average replicative fitness impact of mutations at pleiotropic sites.

### Immune pressure results in multiple co-circulating viral lineages

In the absence of sites impacting antigenicity, our simulations did not reproduce empirically observed patterns of co-circulating viral lineages. However, in the presence of immune pressure, our simulations consistently yielded two, and occasionally three, co-circulating viral lineages (Figures 6N, 6O, 6P; Figures 7M, 7N, 7O, 7P). This was the case for even the lowest extent of immune strength we considered (*q* = 0.3; Figure 6N) and for even relatively broad immunity breadth (*p* = 0.95; Figure 7P). These results make sense in that immune pressure, as implemented, results in negative frequency-dependent selection. As such, the consensus genotype is expected to shift back and forth between the two clades, with the viral clade that does not include the consensus genotype having a selective advantage.

The co-circulation of viral lineages that are observed in the presence of immune pressure also help us understand the seemingly jagged nonsynonymous and synonymous substitution rates apparent in Figures 6F-H and 7E-H. These occur because of co-circulating clades that have different numbers of nonsynonymous as well as synonymous substitutions. The jaggedness appears from the consensus genotype rapidly changing from being in one clade to being in another clade.

### Immune pressure increases the likelihood of observing parallel substitutions

Finally, we examined the impact of immune pressure on the occurrence of parallel mutations across individuals. To compare against simulations that included immune pressure, we first simulated a model with no immune pressure by setting the strength of immune pressure to *q* = 0.0. Consistent with our previous results, mean replicative fitness in each individual increased over the 6 months of simulation (Figure 8A). Next, we calculated the number of parallel high-frequency mutations that were shared across individuals, as we did for Figure 5. We again considered only nonsynonymous sites and set the frequency threshold to 20%. Figure 8F again shows that in the absence of immune pressure (*q* = 0.0), parallel mutations do not readily occur. In contrast, in the presence of immune pressure, parallel mutations are more readily observable (Figure 8G-J). This is particularly the case when the strength of immune pressure is strong and when the breadth of the immune response is moderate (Figure 8J). This is because the rate of antigenic evolution is largest under this parameterization (Figure 8O versus Figures 8K-N).

**Figure 8.**
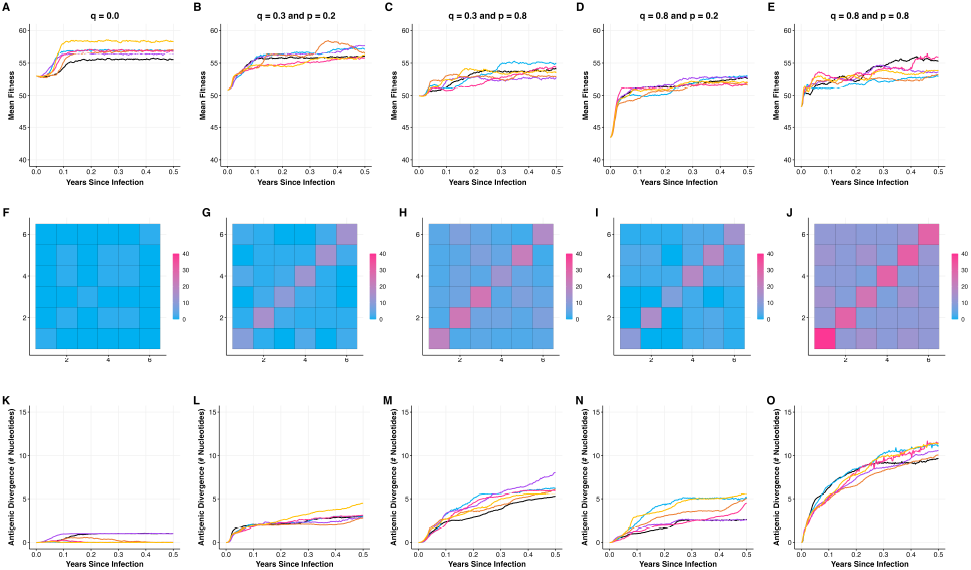
Pleiotropic sites increase the frequency of parallel mutations observed across individuals. Columns correspond to varying parameterizations of the immune response that are also depicted in Figure 1H. Column 1: no immune pressure (*q* = 0.0). Column 2: weak immune strength (*q* = 0.3) and narrow immune breadth (*p* = 0.2). Column 3: weak immune strength (*q* = 0.3) and moderate immune breadth (*p* = 0.8). Column 4: strong immune strength (*q* = 0.8) and narrow immune breadth (*p* = 0.2). Column 5: strong immune strength (*q* = 0.8) and moderate immune breadth (*p* = 0.8). Infecting genotypes are approximately 50% adapted to the host. (A-E) Changes in mean viral replicative fitness for six viral populations evolving on the same fitness landscape, starting with the same infecting genotype. (F-J) The number of shared mutations across pairs of individuals. Only mutations at nonsynonymous sites that exceeded frequencies of 20% at time *t* = 0.5 years were considered in this calculation. (K-O) Extent of antigenic evolution over the course of infection. All simulations were performed using viral genome of length *L* = 400, with *L*_*S*_ = 85, *L*_*P*_ = 267, *L*_*PA*_ = 48, and *L*_*A*_ = 0. Other parameters are: *N* = 5000, *µ* = 2.5 *×* 10^*−*5^ mutations per site per infection cycle, *k* = 100, *c* = 0.2, and *d* = 4 infection cycles per day.

## Discussion

Prolonged infections with respiratory viruses such as influenza viruses and coronaviruses have been extensively documented. Through serial sampling of these infections, several consistent patterns of viral evolution have been identified, including an observed excess of nonsynonymous substitutions (Figure 1A), co-circulating viral lineages (Figure 1B), parallel substitutions across infected individuals (Figure 1C), and variable rates of antigenic evolution (Figure 1D). Here, we have developed a fitness landscape model to determine what processes likely underlie these evolutionary patterns. Through simulation, we found that the patterns shown in Figures 1A-C could not be consistently reproduced in the absence of sites that impacted antigenicity. In contrast, when we instead assumed that a subset of the nonsynonymous sites impacted antigenicity in addition to viral replicative fitness, our simulations were able to consistently reproduce these three observed patterns. The final pattern, of variable rates of antigenic evolution, could be explained by either differing strengths of immune pressure across infected individuals and/or differing breadths of the immune response across infected individuals.

We designed our model to be flexibly parameterized by easily changing the number and types of sites, the mutation rate, the ruggedness of the replicative fitness landscape, and the strength and breadth of the immune response. While we could not simulate the model under all possible parameterizations, we considered the impact of multiple different fitness landscapes on patterns of within-host viral evolution as well as the impact of different strengths and breadths of the immune response on these patterns. Our model did adopt three sets of assumptions that could be relaxed in future studies. First, we assumed that no recombination occurs within our model genome. This assumption simplified our model considerably, in that we did not have to model cellular co-infection either implicitly or explicitly, and it allowed us to organize our sites in the model genome in an arbitrary order. While this assumption simplified our model, it also prevented us from being able to determine how different fitness landscapes and types of immune pressure would impact rates of recombination and ultimately adaptation and immune escape. Second, we assumed that viral population sizes were constant over the course of infections and compared across simulations that had the same viral population size. This assumption could be easily relaxed, but in the absence of a question related specifically to population sizes and population dynamics, we adopted this assumption to be able to interpret our results without this variation as a confounding factor. Third, we assumed a single viral population within each individual, with no spatial subdivision or tissue compartmentalization. As such, spatial structure could not be assessed or invoked as a driver of any of the evolutionary patterns we considered. Given findings that within-host spatial structure occurs in respiratory virus infections and may contribute to genetic diversification of within-host viral populations (Gallagher *et al*., 2018; Chaguza *et al*., 2023; Smith et al., 2024; Farjo et al., 2024; Ferreri et al., 2025), this assumption could be relaxed in future extensions of the model. Finally, we used our model to evaluate the drivers of four different evolutionary patterns that have been documented in prolonged infections with respiratory viruses. As additional studies accrue, some of these patterns themselves may need reevaluation. For example, a recent study has found that sequencing errors may have inflated estimates of viral diversity and evolutionary rates of SARS-CoV-2 populations sampled from individuals experiencing prolonged infections (Rutsinsky *et al*., 2024).

Although the work presented here focused on viral evolution in individuals experiencing prolonged infection with respiratory viruses that generally cause acute infection, our results reveal patterns similar to those observed in other prolonged viral infections, including prolonged infections with enteric viruses such as noroviruses (van Beek *et al*., 2017; Doerflinger et al., 2017) as well as with viruses such as hepatitis C virus (HCV) and HIV that lead to chronic infection. For example, chronic HCV infections are known to result in rapid diversification from the founder virus into multiple distinct viral lineages that continue to cocirculate over the course of the infection (Raghwani *et al*., 2016, 2019). This co-circulation of viral lineages results in significant fluctuations in viral divergence over time (Raghwani *et al*., 2016), a pattern that we also observed in our analysis when we incorporated immune pressure. As such, we hope that the fitness landscape model presented here may further be adapted to evaluate the drivers of observed patterns of viral evolution in these other types of viruses.

## Funding

Research reported in this publication was funded by NIH R01 AI154894 and the National Institute of Allergy and Infectious Diseases, Centers of Excellence for Influenza Research and Response, contract number 75N93021C00017. AC was futher supported by Emory University and the National Institute of Allergy and Infectious Diseases of the National Institutes of Health under Award Number T32AI138952 (Infectious Diseases Across Scales Training Program; IDASTP). The content is solely the responsibility of the authors and does not necessarily represent the official views of the National Institutes of Health.

## Data Availability

All simulation code for the model will be provided on GitHub. All data in this work are simulated data.

## Notes

### Competing Interest Statement

The authors have declared no competing interest.

