## Supplemental Material for "Immune Pressure is Key to Understanding Observed Patterns of Respiratory Virus Evolution in Prolonged Infections"

Amber Coats[1,*], Yintong Rita Wang[2], Katia Koelle[2,3,*]

1 Program in Microbiology and Molecular Genetics, Emory University, Atlanta, GA

2 Department of Biology, Emory University, Atlanta, GA

3 Emory Center of Excellence for Influenza Research and Response (CEIRR), Atlanta GA, USA

*

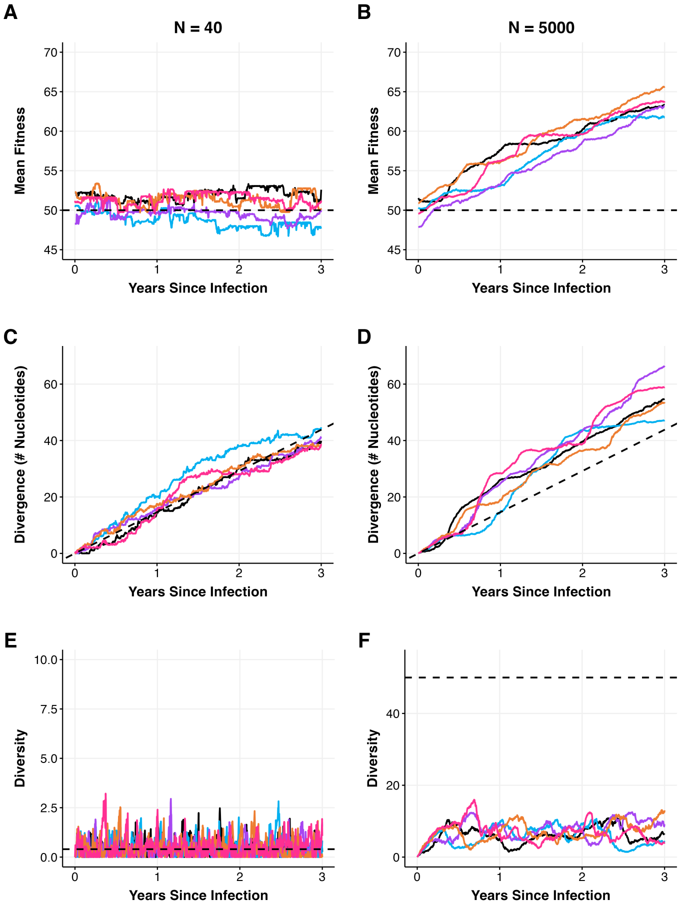


**Figure S1. Genetic drift and selection in within-host viral populations of different size.** The first column shows viral evolutionary dynamics of six independently simulated prolonged infections where the size of the viral population is $N = 40$. The second column shows viral evolutionary dynamics of another six independently simulated prolonged infections where the viral population size is $N = 5000$. (A,B) Mean replicative fitness of evolving viral populations. Dashed black lines show expected fitness of the infecting genotype. (C,D) Mean nucleotide divergence of the populations shown in (A,B). Dashed black lines show expected divergence under neutral evolution, given by $\mu L t$, where $L$ is the genome length, time $t$ is measured in years, and the mutation rate, converted to the time scale of years, is: $\mu = 0.0365$ mutations per site per year ($= 2.5 \times 10^{-5}$ mutations per site per replication cycle $\times 4$ replication cycles per day $\times 365.25$ days per year). (E,F) Mean pairwise diversity of the populations shown in (A,B). The dashed black lines show expected equilibrium diversity levels under neutral evolution. The expected level of pairwise diversity (of a haploid population evolving under a Moran model) is given by: $\mu \times L \times N$. The evolutionary dynamics of all viral populations were simulated for three years. All simulations used a genome length of $L = 400$ nucleotides, with synonymous $L_S = 85$ sites and phenotypic $L_P = 315$ sites ($L_A = 0$, $L_{PA} = 0$). Other parameters are: $c = 0.5$, $\mu = 2.5 \times 10^{-5}$ mutations per site per replication cycle, $k = 100$, $d = 4$ per day, and $\Delta t = 1$ hr. Each infecting genotype had a Hamming distance of exactly 200 nucleotides from the reference genotype $G^*$ of all ones.

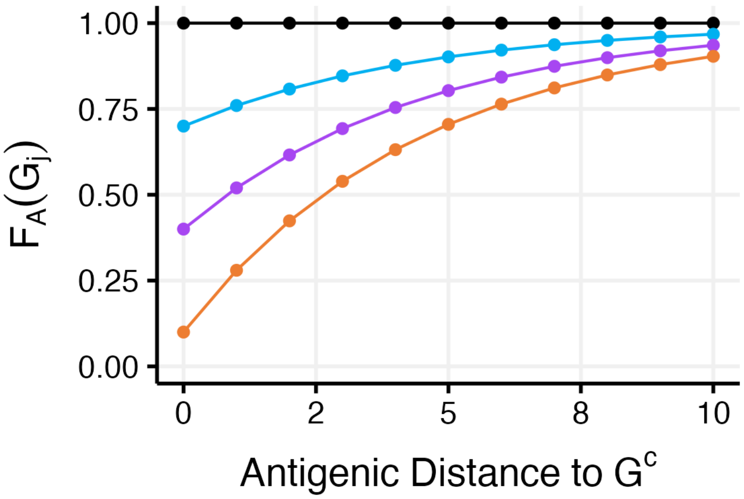


**Figure S2. Antigenic fitness parameterizations when considering the impact of different strengths of immune pressure on viral evolution.** The plot shows the antigenic fitness of genotypes that are different antigenic distances away from the consensus genotype $G^c$. The black line shows the antigenic fitness function parameterized with $q = 0$. The light blue line shows $F_A$ with $q = 0.3$, corresponding to immune pressure being low. The purple line shows $F_A$ with $q = 0.6$, corresponding to immune pressure being moderate. The orange line shows $F_A$ with $q = 0.9$, corresponding to immune pressure being strong. The breadth of the immune response in all cases considered here is set to $p = 0.8$.

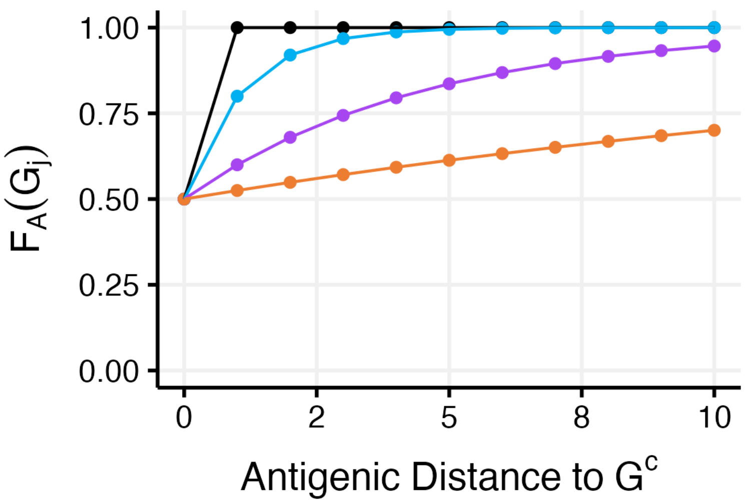


**Figure S3. Antigenic fitness parameterizations when considering the impact of different breadths of the immune response on viral evolution.** The plot shows the antigenic fitness of genotypes that are different antigenic distances away from the consensus genotype $G^c$. The black line shows the antigenic fitness function parameterized with $p = 0.0$. The blue line shows $F_A$ with $p = 0.4$. The purple line shows $F_A$ with $p = 0.8$. The orange line shows $F_A$ with $p = 0.95$. The strength of the immune response in all cases considered here is set to $q = 0.5$.

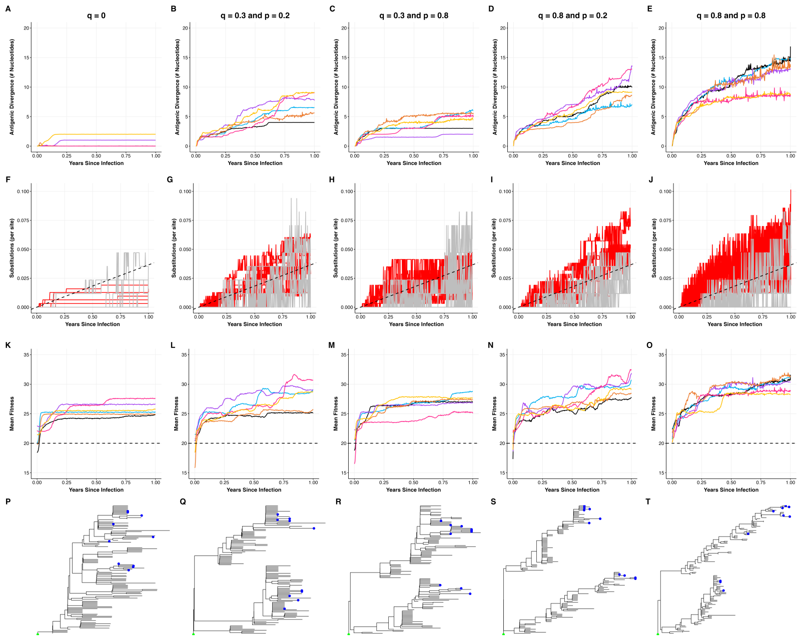


**Figure S4. Immune pressure facilitates viral adaptation when the infecting genotype is poorly adapted to the host.** Columns correspond to varying parameterizations of the immune response that are also depicted in Figure 1H. Column 1: no immune pressure ($q = 0.0$). Column 2: weak immune strength ($q = 0.3$) and narrow immune breadth ($p = 0.2$). Column 3: weak immune strength ($q = 0.3$) and broad immune breadth ($p = 0.8$). Column 4: strong immune strength ($q = 0.8$) and narrow immune breadth ($p = 0.2$). Column 5: strong immune strength ($q = 0.8$) and broad immune breadth ($p = 0.8$). Infecting genotypes are approximately 20% adapted to the host. (A-E) Extent of antigenic evolution over the course of infection for six simulations. Antigenic evolution was calculated as divergence between the consensus genotype at a given time point and the infecting genotype, at the subset of nonsynonymous sites that impacted antigenicity. (F-J) Number of nonsynonymous (red) and synonymous (grey) substitutions per site over the course of infection. Dashed black line shows the expected number of substitutions under neutral evolution. (K-O) Mean viral replicative fitness over the course of infection. The horizontal dashed line shows the expected fitness of the infecting genotype. (P-T) Neighbor-joining trees, generated using 10 randomly sampled genotypes every 30 days. NJ trees are shown for the the simulation shown in black in the above panels. All trees were rooted on the infecting genotype. Simulations were performed using a viral genome of length $L = 400$, with $L_S = 85$, $L_P = 267$, $L_{PA} = 48$, and $L_A = 0$. Other parameters are: $N = 5000$, $\mu = 2.5 \times 10^{-5}$ mutations per site per infection cycle, $k = 100$, $c = 0.2$, and $d = 4$ infection cycles per day.

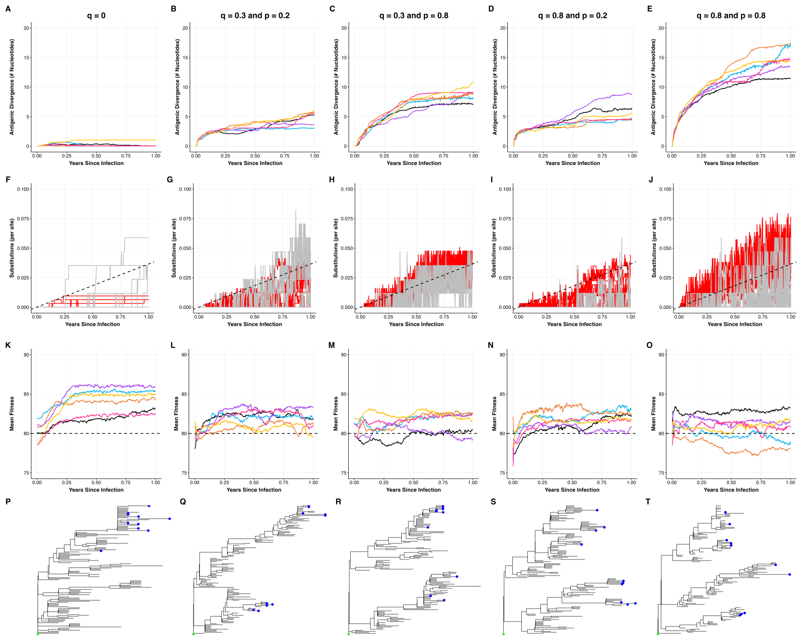


**Figure S5. Immune pressure impedes viral adaptation when the infecting genotype is well adapted to the host.** Columns correspond to varying parameterizations of the immune response that are also depicted in Figure 1H. Column 1: no immune pressure ($q = 0.0$). Column 2: weak immune strength ($q = 0.3$) and narrow immune breadth ($p = 0.2$). Column 3: weak immune strength ($q = 0.3$) and moderate immune breadth ($p = 0.8$). Column 4: strong immune strength ($q = 0.8$) and narrow immune breadth ($p = 0.2$). Column 5: strong immune strength ($q = 0.8$) and moderate immune breadth ($p = 0.8$). Infecting genotypes are approximately 80% adapted to the host. (A-E) Extent of antigenic evolution over the course of infection for six simulations. Antigenic evolution was calculated as divergence between the consensus genotype at a given time point and the infecting genotype, at the subset of nonsynonymous sites that impacted antigenicity. (F-J) Number of nonsynonymous (red) and synonymous (grey) substitutions per site over the course of infection. Dashed black line shows the expected number of substitutions under neutral evolution. (K-O) Mean viral replicative fitness over the course of infection. The horizontal dashed line shows the expected fitness of the infecting genotype. (P-T) Neighbor-joining trees, generated using 10 randomly sampled genotypes every 30 days. NJ trees are shown for the the simulation shown in black in the above panels. All trees were rooted on the infecting genotype. Simulations were performed using a viral genome of length $L = 400$, with $L_S = 85$, $L_P = 267$, $L_{PA} = 48$, and $L_A = 0$. Other parameters are: $N = 5000$, $\mu = 2.5 \times 10^{-5}$ mutations per site per infection cycle, $k = 100$, $c = 0.2$, and $d = 4$ infection cycles per day.
